# RNA tertiary structure modeling with BRiQ potential in CASP15

**DOI:** 10.1101/2023.05.26.542548

**Authors:** Ke Chen, Yaoqi Zhou, Sheng Wang, Peng Xiong

## Abstract

We describe the modeling method for RNA tertiary structures employed by team AIchemy_RNA2 in the 15^th^ Critical Assessment of Protein Structure Prediction (CASP15). The method consists of the following steps. Firstly, secondary structure information was derived from various manually-verified sources. With this information, the full length RNA was fragmented into structural motifs. The structures of each motif were predicted and then assembled into the full structure. To reduce the searching conformational space, a RNA structure was organized into an optimal base folding tree. And to further improve the sampling efficiency, the energy surface was smoothed at high temperatures during the Monte Carlo sampling to make it easier to move across the energy barrier. The statistical potential energy function BRiQ was employed during Monte Carlo energy optimization.

## 1. Introduction

RNA is a special biological macromolecule that can both transfer genetic information and perform biological functions, including catalysis^1,2^ and gene regulation^2,3^. These functions depend on the specific tertiary structure of RNA^4,5^, therefore, characterization of RNA structures is of crucial importance^6^. Among various methods serving for this purpose, computational modeling of RNA structures is developing into an important and rapidly-advancing field^5,6^. Similar to prediction of protein structures, a variety of approaches have been developed for predicting the RNA 3D structure^7^, including homology modeling^8,9^, fragment assembly^10–16^, and *de novo* prediction with coarse grained models^17–20^ as well as deep learning methods^21–26^, following the success of AlphaFold2^27^. Since 2011, blind tests of RNA structure prediction were carried out in RNA-puzzles (https://www.rnapuzzles.org) to evaluate the capabilities and limitations of current methods. The test was introduced to 15^th^ Community Wide Experiment on the Critical Assessment of Techniques for Protein Structure Prediction (CASP15, https://predictioncenter.org/casp15) for the first time and received unprecedented attention.

Our team (AIchemy_RNA2) took part in the RNA 3D structure prediction experiment in CASP15 with the best performance. We followed a classic energy-based strategy to predict RNA structure, whose performance depends on the accuracy of the energy potential, and the effectiveness of conformational sampling. In our previous work, we proposed a Backbone Rotameric and Quantum mechanical energy scaled base-base knowledge-based potential (BRiQ) for RNA structure refinement^28^. Unlike other mostly coarse-grained statistical potentials, the BRiQ potential is a full-atom potential that includes all bonded and non-bonded interactions. In addition, Unlike a traditional pairwise all-atom molecular force field such as AMBER^29^ equipped with analytic formular, the BRiQ potential is high dimensional in discrete numerical representations. To optimize this statistical potential, we developed a base folding tree algorithm to sample the RNA structural space. A number of strategies were employed to reduce the conformational space and increase the efficiency of sampling. The most important strategy can be described as divide and conquer: RNA structure motifs were predicted first and then assembled into the full model for refinement. For some targets, the restraints derived from homology models were used. The overall pipeline is not yet fully automatic. Manual intervention is necessary in most cases.

## 2. Methods

### 2.1 Overview of the pipeline

In CASP15, we participated in the RNA tertiary structure category with group ID AIchemy_RNA2. The overview of the prediction pipeline is shown in Figure 1. Starting from the target sequence, we first performed a Needleman-Wunsch algorithm which align the sequence against the PDB database to detect homology structure. Then, we attempted various strategies to obtain the secondary structure of the target sequence, including homology analysis, literature search and our own energy-based secondary structure prediction program. With reliable secondary structure information, tertiary structure motifs were identified by human intuition and predicted using RNA-BRiQ programs. For each structure motif, we generated thousands of decoy models. These models were clustered using a density based clustering algorithm. Representative models were selected for the assembling procedure. Finally, the overall structures were optimized using the RNA-BRiQ program and five models for each target were selected.

**Figure 1.**
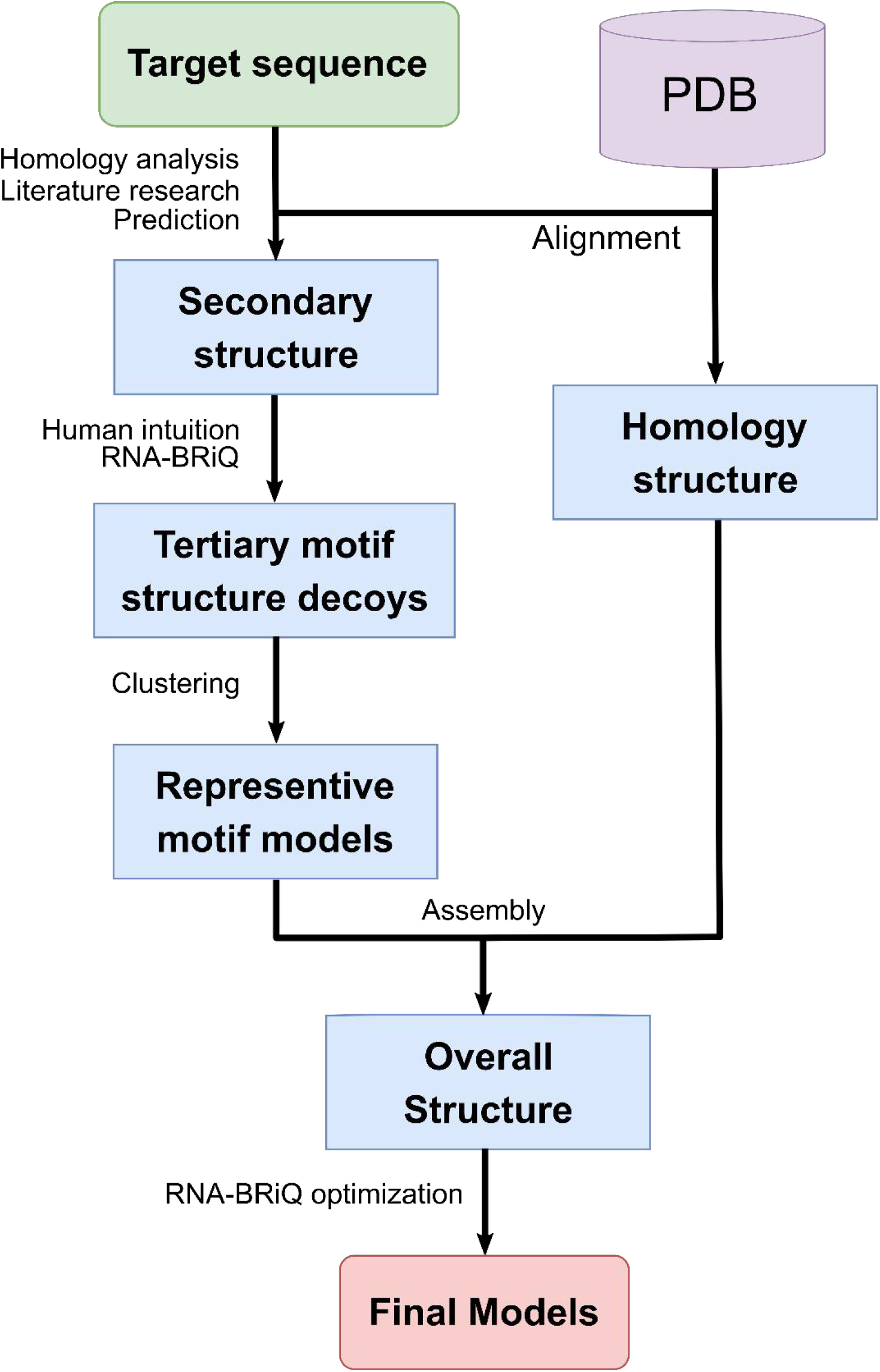
The schematic diagram of the RNA tertiary structure modeling pipeline

### 2.2 Representation of an RNA molecule

Unlike other coarse-grained methods for RNA structure modeling, BRiQ utilizes a full-atom representation, except for hydrogen atoms which are treated as a part of the neighboring heavy atoms. Each residue is composed of three groups: base, ribose and phosphate. The base is considered as a rigid body consisting of 8-11 non-hydrogen atoms. The position of base atoms are described by a local frame (cs_res_) defined by atom C1’, N9 and C4 for adenine and guanine or C1’, N1 and C2 for cytosine and uracil. The ribose contains 8 atoms: O2’, O3’, and C5’ plus 5 atoms in the sugar ring. The coordinates in the local frame cs_res_ of the base were clustered into 1500 conformers, namely ribose rotamers. Thus, the positions of ribose atoms could be described by the local frame cs1 and the ribose rotamer type. Unlike the PDB format, the phosphate group, considered as a rigid body with 4 atoms (O5’, P, OP1 and OP2), is linked to the atom C3’ in our model, so that the position of phosphate atoms can be described by two dihedral angles only: C2’-C3’-O3’-P (ε) and C3’-O3’-P-O5’ (ζ), while the bond lengths and bond angles are fixed. If the PDB format (link via C5’ atom) was followed, three dihedral angles would be required.

### 2.3 BRiQ potential and structure sampling

We employed the BRiQ potential to guide *ab initio* modeling of structure motifs and to refine the assembled full model. The BRiQ potential includes the following interactions: bonded interactions, base pairing and base stacking, hydrogen bonds between backbone oxygen atoms, the interactions between base and backbone oxygen atoms, and steric clashes.

Bonded interactions for base, ribose and phosphate groups were described differently. The base is considered as a rigid body thus, its bonded interaction is ignored. The ribose is represented by rotamers, whose energies are proportional to −log(rot). For the phosphate group, the energy includes a special form of the dihedral angle energy, which takes into account the coupling between the adjacent three dihedral angles. Base pairing and base stacking energies are described as 6D statistical energies with well depths rescaled according to quantum mechanical calculations. Other polar interactions are described by 3D or 4D statistical terms. There is also an energy term for steric clashes (without attractive terms in the van der Waals energy).

To make it easier to move across an energy barrier, we have smoothed the energy landscape. In conformational sampling, the highest energy barrier originates from bond breaking and steric clashes. We employed two strategies to reduce the barrier. First, we replaced the high energy region for chain breaking and steric clashes by a linear extension. During simulated annealing, we imposed weak penalties for bond breaking and steric clashes at high temperature and strong penalties at low temperature. This parameter changes make a more efficient sampling of conformational space.

The BRiQ potential is coupled with a nucleobase-centric tree (NuTree) algorithm for conformational sampling, in which each node denotes a residue and each edge represents the relative position between two local frames of the base, or a SE(3) transformation. The edges in the NuTree connecting Watson-Crick pairs, non-Watson-Crick pairs, sequential stacking neighbors, sequential free neighbors, 1-3 non-sequential stacking neighbors and others were considered as different edge types. Each edge type corresponds to a specific SE(3) transformation space with different size. The space size for sequential free neighbors is much larger than that for Watson-Crick pairs and for sequential stacking neighbors. There is no unique method for constructing a NuTree. In CASP15, many NuTrees were constructed manually to minimize the conformational space.

Furthermore, we have two types of constraints: those acting on the nodes and those acting on the edges. These constraints were applied to fix the structure motifs generated from homologous templates (if any), for example the G-quadruplex in R1126 and R1136, and the helical regions in large RNA molecules to increase sampling efficiency, for example, R1138.

### 2.4 Homology template detection

Firstly, we adopt the Needle-Wunsch algorithm to search the target sequence against the PDB sequence library. For target R1107 and R1108, we find a hit 4pr6 with sequence identity of 62.3%, which was employed as the structure template. For target R1116, template 7lyg can match to 30 nucleotides on 5’ end and 37 nucleotides on 3’ end. As a result, we only predicted the central regions. For target R1117, we employed template 3fu2 with a sequence identity of 50% and consistent secondary structure. For synthetic RNAs, we found no templates but detected ligand binding motifs. Target R1126 contains a 1TU binding motif, whose template is 4kzd. Target R1136 contains a 1TU binding motif and a J93 binding motif, whose templates are 4kzd and 7eop. Finally, the template for protein-RNA complex R1189 and R1190 is 2mf0. The ligand name helped us to locate the correct ligand binding motif.

### 2.5 Secondary structure determination

In order to achieve best modeling performance, we made our best effort to employ correct and complete secondary structure as input. For target R1107, R1108, R1116, R1117, R1189 and R1190, secondary structure information could be derived from their corresponding templates. For synthetic RNAs, UUCG tetra loops were utilized at the end of the helixes, which became a cue to infer the secondary structure. For target R1149, R1156 and the middle region of R1116, the secondary structure came from the literature^30–32^, which reported chemical probing results.

### 2.6 Motif assignment and prediction

We employed a simple strategy to assign motifs according to the secondary structure. For a continuous helix, we retrained two base pairs at each end and eliminated the middle region, so that the whole structure was split into discontinuous motifs. For some special cases, ligand binding motifs in targets R1126 and R1136 were identified from the corresponding template, and kissing loops in targets R1126 and R1138 were identified manually. Using the RNA-BRiQ program, we generated about 2000 models for each motif and then selected top 20% low energy models, which were then submitted to the structure clustering program. Here, we employed one example to illustrate the details of motif structure modeling.

The most frequent motif in CASP15 is the helix hinge motif, which appears in target R1126, R1128, R1136, R1138, R1149 and R1156. It’s a recurrent motif in native structures and is a key unit for constructing synthetic RNAs. The sequences of this motif vary in different targets, but they have the same secondary structure and similar 3D structures. Before the start of Monte Carlo sampling, a NuTree is constructed first. The topology of NuTree is not unique, and different NuTrees tend to sample a different structural space. For example, in Figure 2, we constructed two different NuTrees for the helix hinge motif. For each NuTree, we generated 2000 decoy models. These models can be grouped into four clusters, models generated by NuTreeA tend to fold into conformation 1 and models generated by NuTreeB tend to fold into conformation 3. For other more complex motifs (e.g., the kissing loop in target R1136 and R1138), we construct the NuTree manually. Proper NuTrees are critical for improving sampling efficiency.

**Figure 2.**
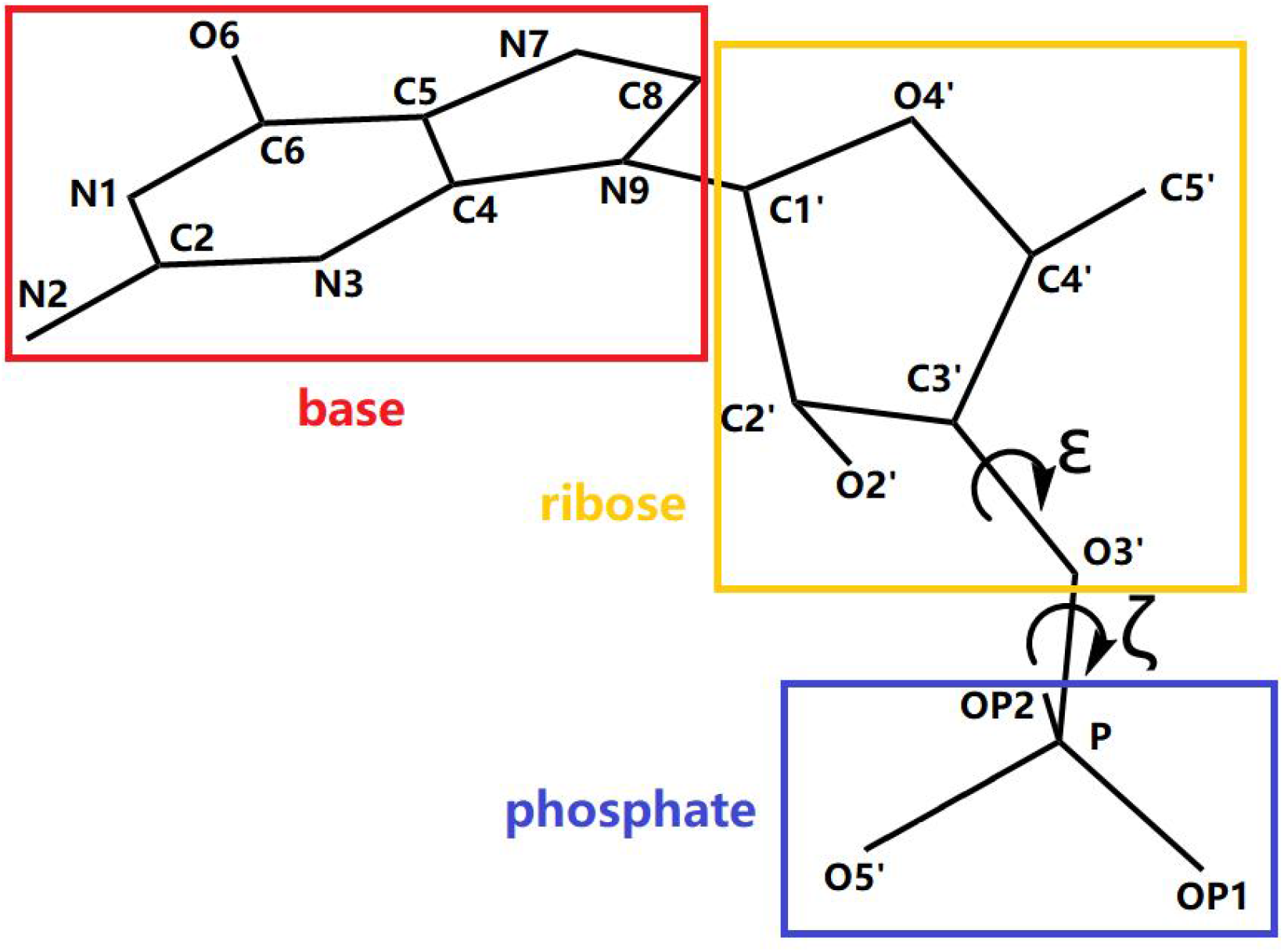
Representation of an RNA residue, the phosphate group is linked to atom O3’ of ribose

**Figure 3.**
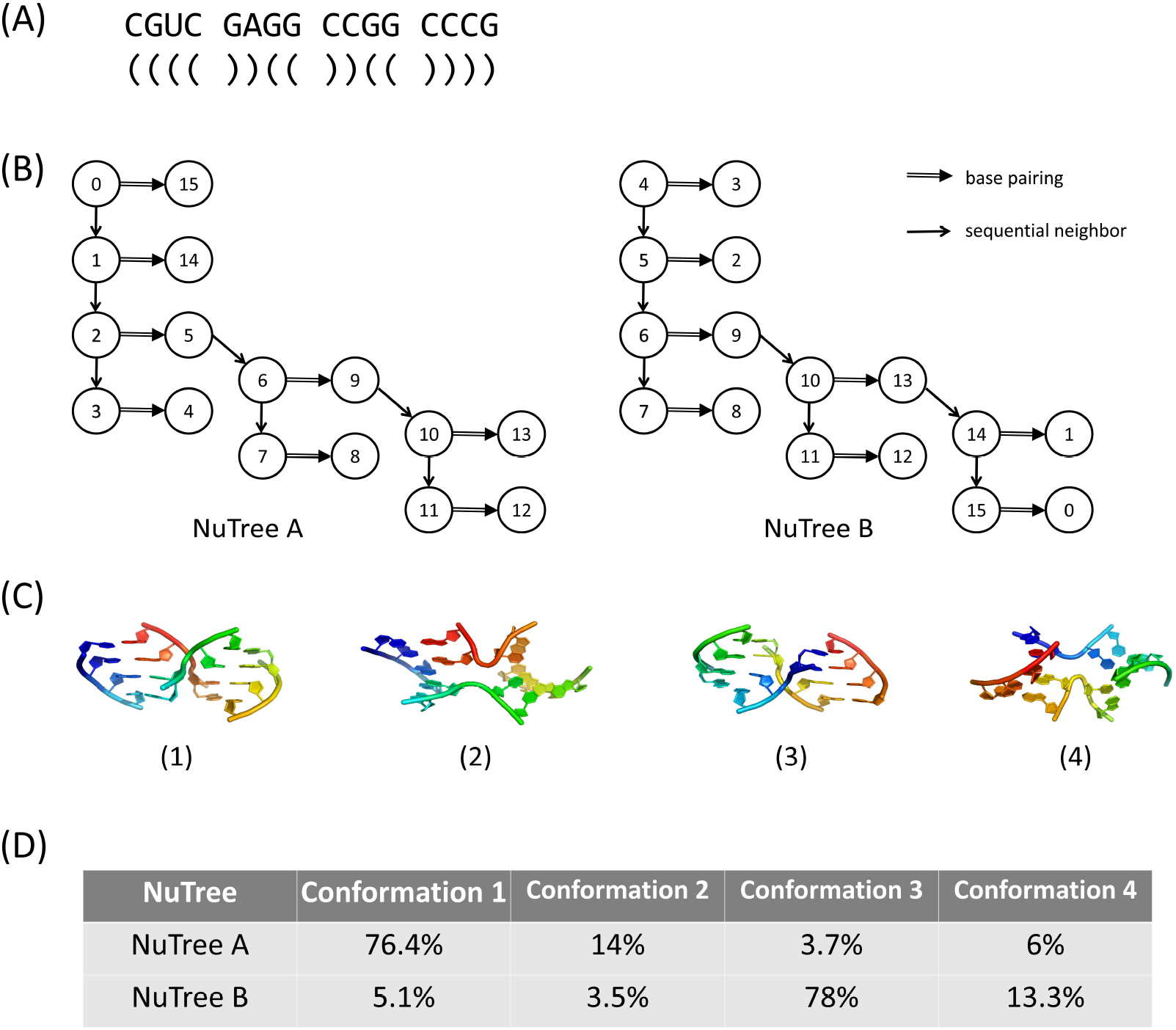
An example of assignment and prediction of the helix hinge motif. (A) The example sequence and secondary structure of the motif. (B) The topologies of two different NuTrees, NuTreeA and NuTreeB for the same sequence. (C) The representative conformations of the four clusters modeled from the NuTrees. These clusters are named after their numbering. (D) The population percentage of each cluster modeled from NuTreeA and NuTreeB, respectively.

### 2.7 Motif assembling and refinement

The time complexity of our sampling algorithm is O(n^3). The computing time rises sharply when the sequence length increases above 50 nucleotides. Therefore, we splited the complete molecule into motifs, predicted the structure of motifs and performed assembling. The boundaries of the motifs are in the helix region. Thus, we made the assembling by aligning the local frame of the residues in helix. When the motif has multiple conformations, such as the four states of the helix hinge motif discussed above, we iterated over all possible combinations and generated different starting models for final refinement. For those large RNAs with multiple motifs, the assembly process is hierarchical, small motifs assembled to large fragments and large fragments assembled to the complete model.

We employed the BRiQ program for final refinement of the assembled structures. In addition to the standard refinement protocol, we included additional constraints to reduce the searching space. There were two types of constraints, those acting on the nodes to fix global coordinates and those acting on the edges to fix relative orientations. In target R1107, R1108 and R1116, the regions that match the structure templates were fixed. In target R1107, R1126 and R1136, the ligand binding motifs were fixed.

## 3. Results and discussion

### 3.1 Overall ranking

When we ranked the performance of different methods based on combined Z-scores, our method is in the leading position. Among the 12 RNA targets in CASP15, six targets (R1107, R1108, R1126, R1128, R1136 and R1138) were ranked first based on the root-mean-squared distance (RMSD). Two additional targets R1117 and R1156 were only slightly worse than the best models from Chen’s group and GeneSilico, respectively. We capture the correct topology of target R1149, but RMSD was a little higher than other groups. Targets R1189 and R1190 were protein-RNA complexes. All models were beyond 15Å RMSD and, thus, the performance ranking for these two targets is not suitable. We did not perform well on targets R1116 as discussed below.

### 3.2 Model accuracy

Here, we go through all targets to review what went right and what went wrong.

For target R1107 and R1108, the X-Ray crystal structures of these two RNAs are homo-dimers, with two monomers connected by four base pairs at position 23-26. Our prediction did not restore the correct dimer state, base 22C and base 37G formed a wrong base pair, causing the conformation of this hairpin loop to deviate from the natural state.

For target R1116, there was a large deviation between our predicted model and the X-Ray crystal structure, the deviation came from the difference between alternative conformations of the helix hinge motif. When we made this prediction, we did not realize that structure sampling results could be dominated by the NuTree topology. As a result, the default NuTree led us to the wrong conformation.

For target R1117, the model we predicted is very close to the natural structure, with an RMSD of 2.27Å, but one base (the cytosine at position 12) was not correctly predicted. In the native structure, the 12C forms a hydrogen bond to O6 atom of guanine at position 3, while in our predicted model, the 12C formed a hydrogen bond to the ribose O2’ atom of position 29. This error caused our predicted model to be slightly worse than that of Chen’s group.

For the four synthetic RNAs, there was no obvious error in our predictions. The RMSDs between predicted models and native structures were ranged from 4.3Å to 8.7Å. Our prediction accuracy was much higher than those of all other groups.

For target R1149 and R1156, the topologies were determined by the helix hinge motif. This time we constructed different kinds of NuTrees manually, and tested all combinations of motif conformations. Our energy function could not tell the difference between these decoys, but at least one of them match to the native structure well with an RMSD of 10.5Å and 7.6Å.

Currently, our program cannot handle the complex structure of protein and RNA, target R1189 and R1190 were modeled poorly.

**Figure 4.**
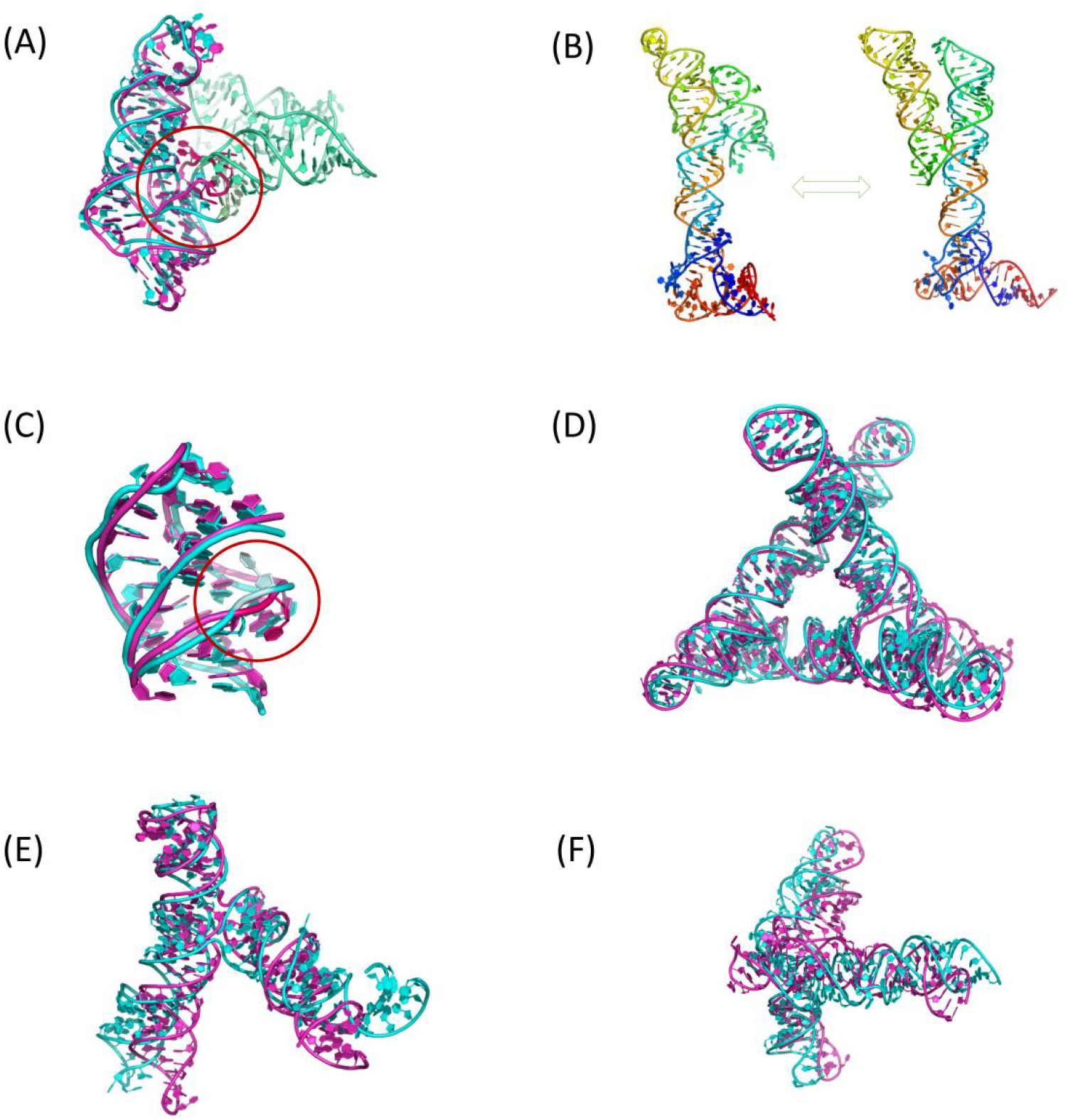
Comparison of native structure and our predicted model, the native structure was shown in cyan or on the left, predicted model was shown in magenta or on the right. (A) Target R1107, the hairpin loop related to dimerization was not predicted well. (B) Target R1116, the topology of helix hinge was wrong. (C) Target R1117, the orientation of cytosine at position 12 was wrong in our predicted model. (D) Target R1128, this target was well predicted. (E) Target R1149, one of our predicted model match to the native structure with RMSD of 10.5Å (F) Target R1156, the native structure has multiple conformations, this is the best aligned prediction, with RMSD of 7.6Å.

## 4. Conclusions

In this work, we have described our pipeline to model the RNA tertiary structures in CASP15. The performance of the pipeline relies on i) the reduction of conformational space brought by the hierarchical modeling framework and proper NuTrees built for motifs, so that it was possible to reach a correct topology in limited time; ii) the high accuracy of the BRiQ potential, which provided advantage especially when target topologies were predicted well by multiple groups. However, the pipeline has not been fully automated that human intuition still plays an important role throughout the pipeline. Additionally, the BRiQ potential is currently inadequate for modeling of protein-RNA complexes. These shortages are awaiting further compliments and we still have a long way to pave for a universal high-resolution modeling method for RNA tertiary structures.

## Acknowledgement

We would like to thank Limin Sheng, Liangzhen Zheng, Minzhi Lin for the support to the project. YZ gratefully acknowledges that the High Performance Computing Cluster at Shenzhen Bay Laboratory was involved in completing this research. He also would like to thank the support of National Key Research and Development Program of China(NO.2021YFF1200400) and the Major Program of Shenzhen Bay Laboratory S201101001, and Shenzhen Science and Technology Program [KQTD20170330155106581].

## Conflicts of interest

The authors declare no competing interests.

## Notes

### Competing Interest Statement

The authors have declared no competing interest.

